# Following The Evolutionary Paths Of Highly Specific Homophilic Adhesion Proteins

**DOI:** 10.1101/2024.03.18.585463

**Authors:** Gil Wiseglass, Rotem Rubinstein

**Affiliations:** School of Neurobiology, Biochemistry and Biophysics, The George S. Wise Faculty of Life Sciences, Tel Aviv University, Tel Aviv, Israel; Sagol School of Neuroscience, Tel Aviv University, Tel Aviv, Israel

## Abstract

Many adhesion proteins, evolutionarily related through gene duplication, exhibit distinct and precise interaction preferences and affinities crucial for cell patterning. Yet, the evolutionary path by which these proteins, which are highly similar in structure and sequence, acquire new specificity and prevent cross-interactions within their family members remains unknown. To bridge this gap, this study focuses on Drosophila Down syndrome cell adhesion molecule-1 (Dscam1) proteins, which are cell adhesion proteins that have undergone extensive gene duplication. Dscam1 evolved under strong selective pressure to achieve strict homophilic recognition, essential for neuronal self-avoidance and patterning. Through a combination of phylogenetic analysis, ancestral sequence reconstruction, and cell aggregation assays, we studied the evolutionary trajectory of Dscam1 exon 4 across various insect lineages. We demonstrated that recent Dscam1 duplications in the mosquito lineage bind with strict homophilic specificities without any cross-interactions. We found that ancestral and intermediate Dscam1 isoforms were able to maintain their homophilic bindings capabilities, with some intermediate isoforms also engaging in promiscuous interactions with other paralogs. Our results highlight the robust selective pressure for homophilic specificity integral to Dscam1 function within the process of neuronal self-avoidance. Importantly, our study suggests that the path to achieving such selective specificity does not introduce disruptive mutations that prevent self-binding but includes an evolutionary intermediate that demonstrates promiscuous heterophilic interactions. Overall, these results offer insights into evolutionary strategies that underlie adhesion protein interaction specificity.

## Introduction

Adhesion proteins play a central role in cellular organization and communication within multicellular organisms (Dalva, et al. 2007; Makrilia, et al. 2009; Lele and Hindges 2023). These are often members of large protein families that have expanded through gene duplication and subsequent evolutionary divergence (i.e., paralog proteins). Due to their ancestral link, paralog proteins generally have similar sequences and nearly identical structures with a tendency for intra-familial interactions (Ispolatov, et al. 2005; Lukatsky, et al. 2007; Pereira-Leal, et al. 2007). Yet adhesion proteins within the same family often display distinct binding specificities and affinities. These differences are central to their roles in cell patterning and organization. Some such roles include neural tube formation mediated by N- and E-cadherins (Taneyhill and Schiffmacher 2017), as well as the function of nectins within the inner ear cell patterning (Togashi, et al. 2011), and clustered protocadherin and Dscam1 proteins in dendritic arborization (Zipursky and Sanes 2010; Honig and Shapiro 2020).

New protein-protein interaction specificities can evolve through mutations that impose negative constraints and prevent cross-interactions among family members (Zarrinpar, et al. 2003; Reinke, et al. 2013; Peleg, et al. 2014; Cheng, et al. 2019; Honig and Shapiro 2020; Sergeeva, et al. 2020). There are two contrasting models that outline the evolutionary pathways proteins undergo from being identical duplicated copies to becoming paralogs with distinct binding specificities. The first model states that over extensive periods of time, proteins lose their functionality due to mutations that cause incompatible interacting interfaces, until additional mutations lead to new interacting interface and binding specificities (Ohno 2013; Siddiq, et al. 2017; McClune and Laub 2020). The second model suggests a continuous evolutionary shift, with intermediate proteins exhibiting promiscuous, non-specific, interactions (Sayou, et al. 2014; Aakre, et al. 2015). While the evolution of specific protein-protein interactions has been extensively studied within the context of enzyme-substrate specificities (Aharoni, et al. 2005; Weinreich, et al. 2006; Khersonsky and Tawfik 2010) and receptor-ligand systems (Chockalingam, et al. 2005; Ortlund, et al. 2007; Eick, et al. 2012; Koehbach, et al. 2013), it remains underexplored from the perspective of cell adhesion proteins.

The Drosophila Down syndrome cell adhesion molecule 1 (Dscam1) serves as an extraordinary example of a large paralogous protein family with highly precise cell surface adhesion interactions. The *Dscam1* gene consists of 24 exons, three of which have undergone extensive duplications (Figure 1A). In alternative splicing, a single exon from each of the three clusters is stochastically selected, with the potential to encode an astounding array of 19,008 unique extracellular regions (Figure 1B). Each extracellular region is characterized by distinct combinations of three alternative immunoglobulin (Ig) domains (Schmucker, et al. 2000).

**Figure 1.**
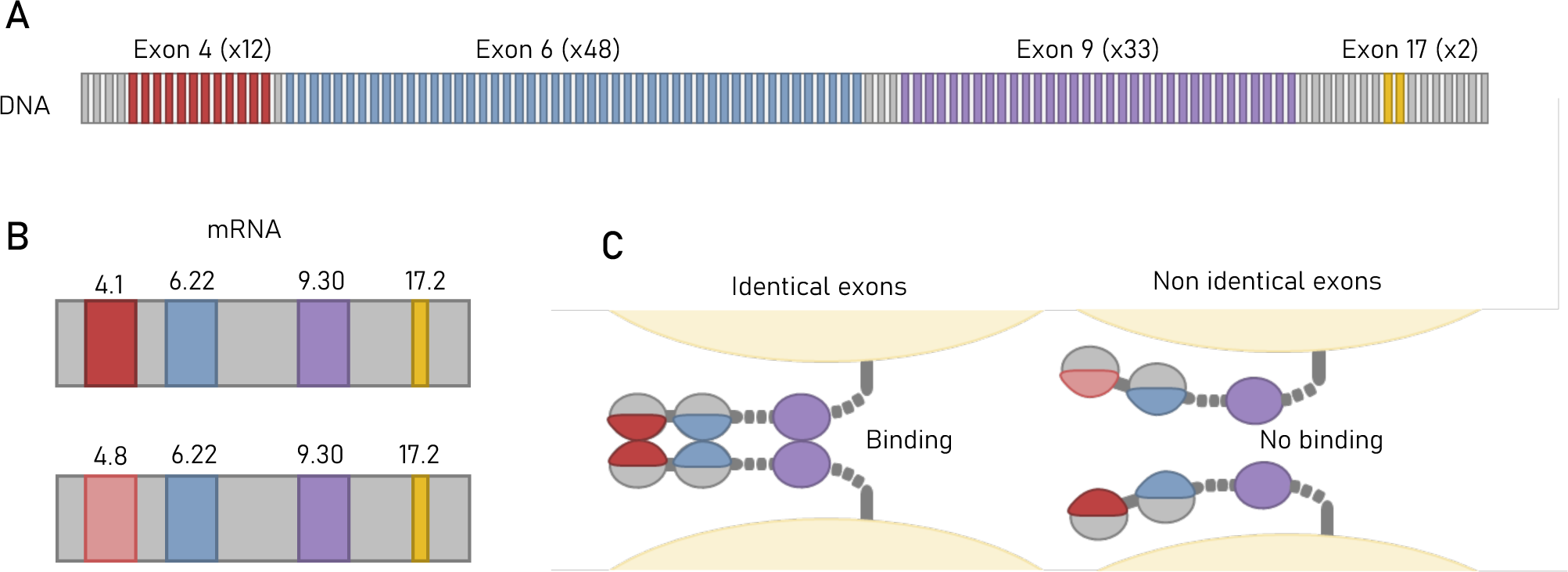
Dscam1 gene has the capacity to transcribe tens of thousands of strictly homophilic isoforms. (A) In insects, exons 4, 6 and 9 of the Dscam1 gene have undergone extensive tandem duplications, contributing to the vast diversity of isoforms produced by the gene. (B) Through stochastic alternative splicing, a single exon from each cluster is retained in the mature transcript. In Drosophila, 38,016 unique isoforms can be expressed, with 19,008 unique extracellular domains. Each neuron expresses a different set of Dscam1 isoforms. (C) Exons 4, 6 and 9 encode into partial (exons 4 and 6) or entire (exon 9) immunoglobulin domain. These domains determine the binding specificity on the Dscam1 protein. For a Dscam1 dimer to form between two cell membranes, all domains must fully match.

Dscam1 isoforms are also unique due to their strict homophilic binding specificity (Figure 1C) (Wojtowicz, et al. 2004; Wojtowicz, et al. 2007). This is in contrast to other cell adhesion protein families that typically exhibit homophilic and heterophilic binding between members of the same family (Katsamba, et al. 2009; Vendome, et al. 2014; Mosca 2015; Zinn and Özkan 2017; Brasch, et al. 2018; Honig and Shapiro 2020; Sergeeva, et al. 2020). The strict homophilic binding exhibited by Dscam1 enormous isoform repertoire is key for its ability to differentiate self from non-self cell-cell interactions. This ability is required for neural patterning within the developing Drosophila nervous system (Wang, et al. 2002; Zhan, et al. 2004; Zhu, et al. 2006; Hughes, et al. 2007; Matthews, et al. 2007; Soba, et al. 2007; Wojtowicz, et al. 2007; Wu, et al. 2012; Miura, et al. 2013; Wilhelm, et al. 2022).

Earlier investigations of Dscam1 evolution traced the extensive exon expansion to the last common ancestor of the Pancrustacea (Lee, et al. 2010; Armitage, et al. 2012), which diverged into insects and crustaceans approximately 500 million years ago (Misof, et al. 2014). These studies used a sample of representative genomes and identified a significant variability in the count of exon duplications among different species, indicating duplication events also occurred in subsequent evolutionary lineages (Lee, et al. 2010; Armitage, et al. 2012). However, these studies did not experimentally test whether newly duplicated Dscam1 isoforms, aside from Drosophila, bind strictly homophilically. The functionality of Dscam1 ancestral proteins also remains unknown, resulting in a knowledge gap in the evolution of adhesion specificity for this unique protein family.

Here, we extensively investigate the evolutionary expansion of Dscam1 exon 4 in insects. We predict the mutational pathways that connect ancestral to recent Dscam1 duplications. Our analysis reveals a clear pattern: as Dscam1 paralogous exons diverge from identical duplicates, some initially exhibit heterophilic interactions, which diminish with additional mutations, ultimately resulting in exclusive homophilic binding. We did not observe a “non-functional” ancestral isoform lacking homophilic binding. By demonstrating this evolutionary progression, we shed light on the key process of increasing the number of highly specific adhesion molecules, deepening our understanding of this critical aspect of molecular evolution.

## Results

### Phylogenetic analysis of Dscam1 exon 4 in insects

In this study, we investigated the evolutionary trajectory of the *Dscam1* gene, focusing specifically on exon 4, which encodes a segment of the Ig2 domain that is involved in homophilic dimerization. A tblastn search was performed against the RefSeq genome database (Johnson, et al. 2008; Coordinators 2016; O’Leary, et al. 2016) for sequences with homology to *Drosophila melanogaster* exon 4.7 (Figure 2A). This comprehensive search covered 83 insect species, resulting in the identification of 962 homologous sequences. A non-redundant set of these sequences, including a Dscam2 sequence from Chelicerata *Ixodes scapularis* as an outgroup, was aligned and used to construct a phylogenetic tree (for additional details, see methods). The resulting tree revealed nine distinct clusters of exons, each representing orthologous sequences from various species (Figure 2B). These orthologous sequences maintain high levels of sequence conservation, particularly in the Ig2 dimer interface, with an average similarity of over 90% within each ortholog cluster. Notably, we found that all clusters contain representative sequences from most species, indicating paralogous relationships between clusters (Figure 2B). For example, the beetle exon 4 sequences can be found in all nine clusters. While previous work suggests that the last common ancestor of insects possessed nine exon 4 paralogs (Lee, et al. 2010; Armitage, et al. 2012), our current findings provide strong support for this notion, mainly due to the increased availability of genomic data.

**Figure 2.**
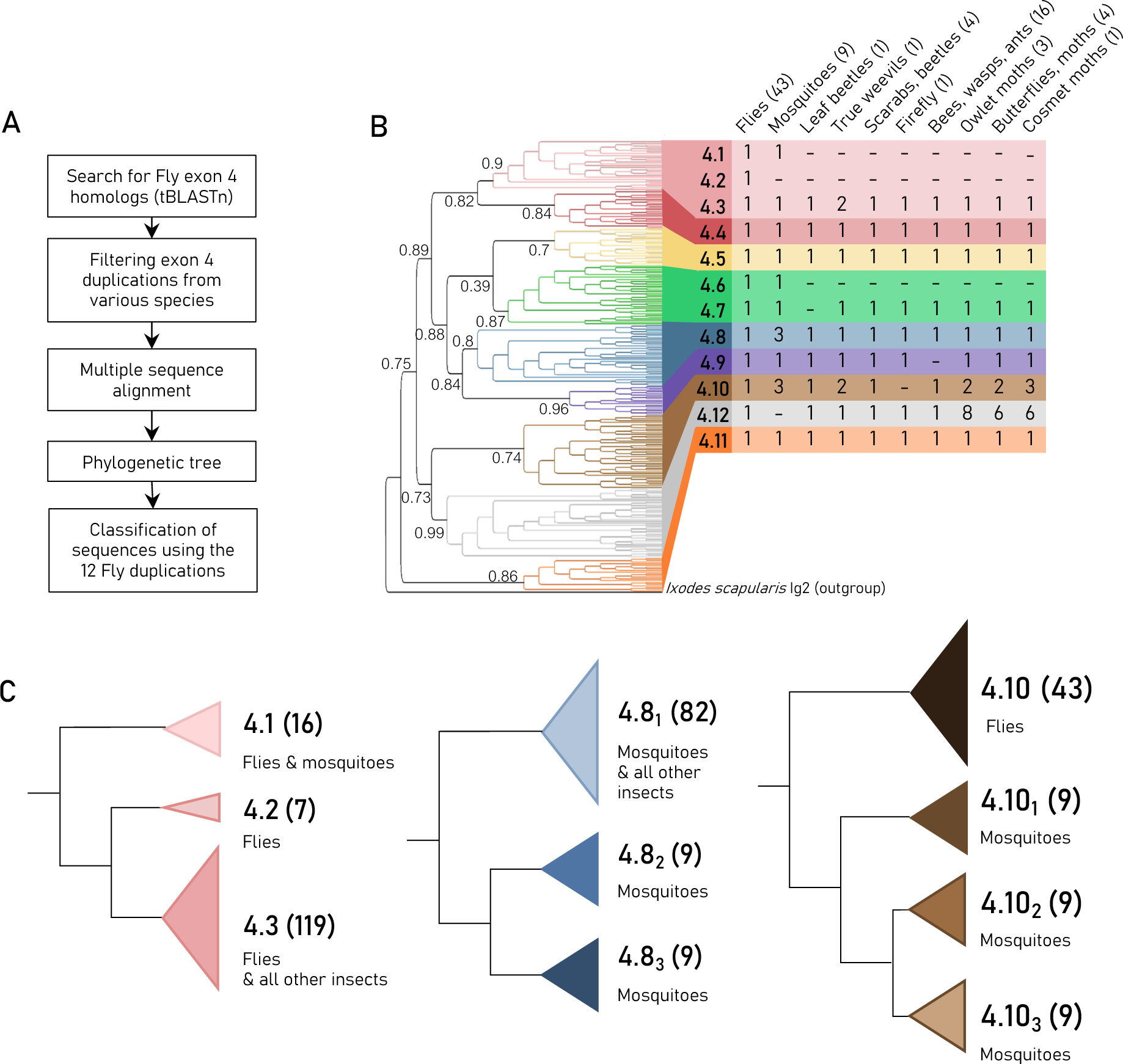
Phylogenetic analysis of insect Exons 4. (A) Illustration of the workflow for the phylogenetic analysis. (B) The phylogenetic tree constructed from 266 exon 4 sequences from 83 representative species (left). The tree topology preserves the organization of the major ortholog clusters, which are highlighted by different colors and are notated according to the Drosophila exon 4 nomenclature. The tree was generated using FastTree with FastTree Local support values shown. The table summarizes the number of duplications (paralogs) per species group for each cluster (right). The number of species per group is denoted within brackets. (C) Simplified phylogenetic sub-trees of three recent duplications (4.1-4.3, 4.81-3, and 4.101-3) for both flies and mosquitoes. The number of sequences per cluster is denoted within the brackets. This number includes the redundant sequences which were not used in reconstructing the main tree. Colors correspond to the main tree and the number of sequences in each clade is indicated in parentheses.

Next, using the 12 exon 4 paralogs of *Drosophila melanogaster* (4.1 to 4.12) as a reference, we focused on more recent exon 4 duplication events occurring in specific insect lineages. These analyses uncovered the absence of Drosophila exons 4.1, 4.2, and 4.6 in most insect species. These exons are unique to the Diptera lineage and encompass flies and mosquitoes, which diverged from other insects approximately 260 million years ago (Wiegmann, et al. 2011). Sequence homology and branch support values strongly indicate recent duplications and divergence of exons 4.1 and 4.2 from exon 4.3 (Figure 2C), as well as the divergence of exon 4.6 from exon 4.7 (Figure 2B). We also discovered more recent lineage-specific duplications, including duplications of exons 4.8 and 4.10 found in mosquitoes (Figures 2C and 3) and duplications of exons 4.10 and 4.12 in the Lepidoptera (i.e., moths and butterflies) (Figure 2B).

**Figure 3.**
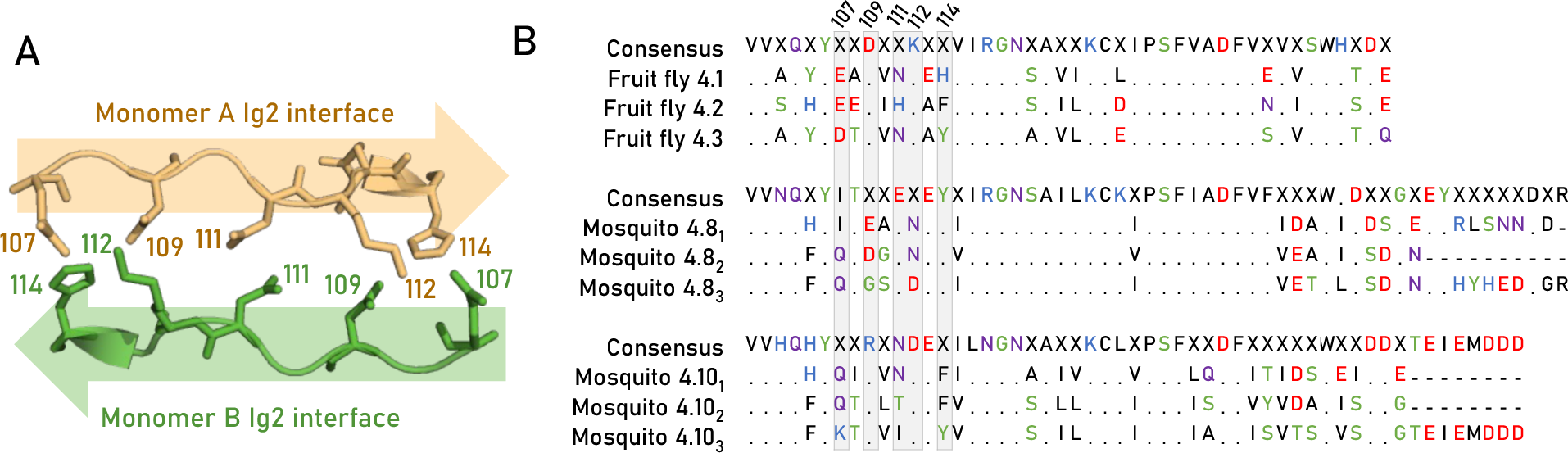
Three recent duplications for both mosquitos and flies. A) structure of the Ig2 homodimer interface (PDB 3DMK) encoded by Drosophila exon. 4.1. The interface spans amino acid positions 107-114 and aligns in an anti-parallel fashion. B) Multiple sequence alignments of the Fruit fly *Drosophila melanogaster* exons 4.1-3 and the Yellow fever mosquito *Aedes aegypti* exons 4.81-3 and 4.101-3. A period (i.e., “.”) in the multiple sequence alignments indicates invariant positions and only residues that deviate from the consensus sequence are shown. The Ig2:Ig2 interface residues are highlighted in grey, and the sequence position is indicated at the top of the alignment.

### Recent duplications in mosquito Dscam1 exhibit strict homophilic binding

To date, the highly specific homophilic dimerization of the Dscam1 protein has been observed exclusively between *Drosophila melanogaster* isoforms. This study aimed to investigate whether the binding specificities of recent exon 4 duplications, not present in Drosophila, would also maintain homophilic specificity. We focused on mosquito exons 4.8 and 4.10, along with their respective duplications, referred to here as 4.8_1_, 4.8_2_, 4.8_3_, 4.10_1_, 4.10_2_, and 4.10_3_. To assess whether the newly duplicated isoforms evolved new homophilic binding specificities, we used site-directed mutagenesis to swap the interface residues of Drosophila Dscam1 to match those found in the mosquito isoforms (Figure 3). We implemented this strategy based on a previous study showing that isoform specificity could be altered by the substitution of residues in positions 107 to 114 on the Ig2 domain dimer interface (Wojtowicz, et al. 2007).

We assessed the binding preferences of the mosquito interfaces via cell aggregation assays using HEK293F-suspended cells. Cell aggregation is a well-established method used to determine the binding specificity of adhesion proteins (Matthews, et al. 2007; Schreiner and Weiner 2010; Boucard, et al. 2014; Thu, et al. 2014; Rubinstein, et al. 2015; Bisogni, et al. 2018; Zhou, et al. 2020; Hou, et al. 2022; Cheng, et al. 2023; Wiseglass, et al. 2023). Each protein is tagged with either red or green fluorescent markers and transfected into separate cell populations. The two cell populations are then mixed and allowed to aggregate based on the binding specificities of the adhesion proteins they express. If the two proteins are strictly homophilic, the cells will form separate red or green aggregates. In contrast, if the two proteins are heterophilic, the cells will form mixed red and green aggregates. To quantify the extent of aggregate mixing or separation, we employed a customized Python script (Wiseglass, et al. 2023), calculating the ratio of proximate red and green cells (see methods). The resulting ratio is displayed in the corner of each image, with a value exceeding 0.1 indicating visibly mixed aggregates (Figure 4).

**Figure 4.**
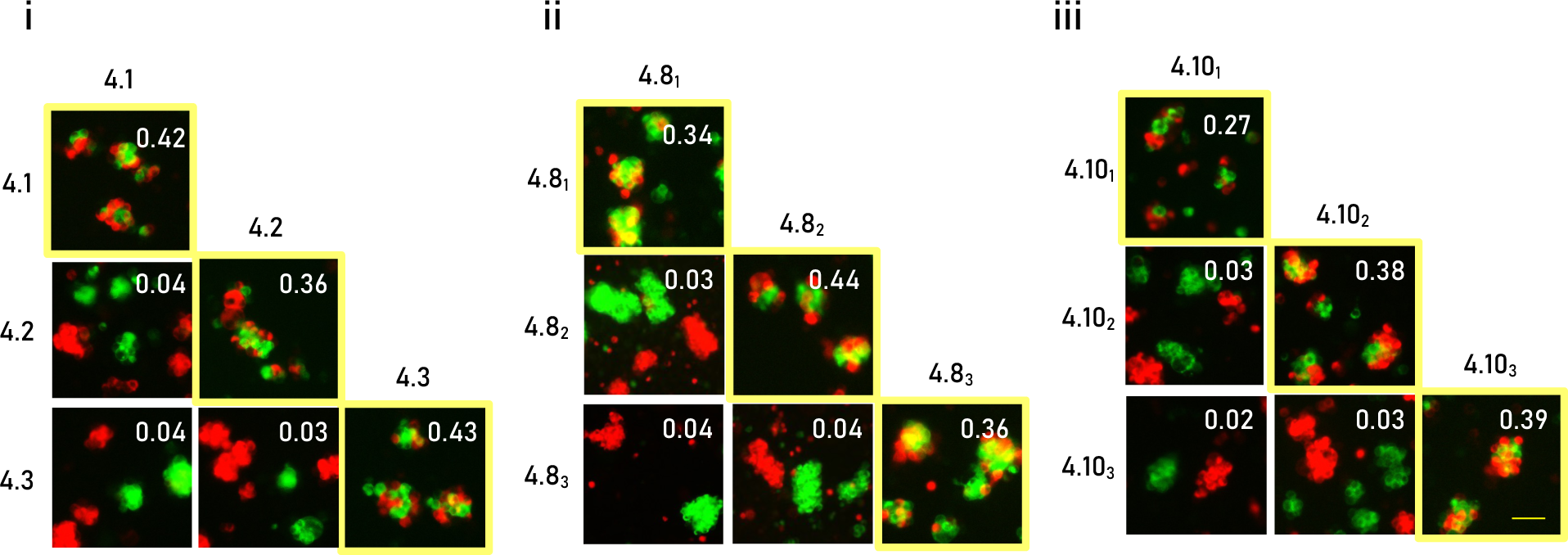
Recent duplications in mosquito Dscam1 engage in highly specific homophilic interactions. Pairwise combinations within each exon paralog cluster were assessed for their interaction specificity. HEK293F cells expressing identical isoforms formed mixed red and green aggregates, while cells expressing different isoforms formed separate red and green aggregates. i) Binding assay for the Fruit fly *Drosophila melanogaster* exons 4.1-3. ii and iii) Binding assays for the Yellow fever mosquito *Aedes aegypti* exons 4.8_1-3_ and 4.10_1-3_, respectively. The aggregate mixing score is presented in the right corner of each image. Scale 100 µm.

Cell aggregation assay was performed in pairwise combinations of the yellow fever mosquito *Aedes aegypti* isoforms 4.8_1_-4.8_3_, 4.10_1_-4.10_3_, and the *Drosophila melanogaster* isoforms 4.1-We observed that only cells expressing identical isoforms formed mixed aggregates, while all combinations of non-identical isoform pairs resulted in separate aggregates (Figure 4). These results demonstrate a strict homophilic binding preference for each isoform. Importantly, our findings indicate that the recent mosquito exons evolved to encode adhesion receptors with highly specific homophilic cell recognition. This suggests mosquito Dscam1 has a similar function to Drosophila Dscam1 in mediating the distinction between self and non-self in neurons.

### Tracing evolutionary paths of binary specificities in Dscam1

The evolution of Dscam1 provides a unique opportunity to explore the challenges associated with diversifying homophilic interfaces. Following exon duplication, alterations in the dimer interface can ultimately lead to the establishment of a new homophilic specificity. However, during this evolutionary process, intermediate changes may result in a non-specific binding or potential loss of binding altogether. To gain deeper insights into the evolutionary trajectory of Dscam1 and to predict intermediate isoforms, we performed an ancestral sequence reconstruction. Utilizing two ancestry sequence reconstruction programs, PaML (Yang 2007; Xu and Yang 2013) and GRASP (Ross, et al. 2022), we predicted the last common ancestor (LCA) proteins prior to the recent mosquito duplication of exons 4.8 and 4.10, and the duplication in Drosophila exon 4.3, referred to here as, 4.8_LCA_, 4.10_LCA_, and 4.1-3_LCA_. The reconstruction of the three ancestor proteins achieved high confidence of 0.85, 0.91, and 0.84 (for 4.1-3_LCA_, 4.8_LCA_, and 4.10_LCA_ respectively) with predicted ancestral interface residues similar, but not identical, to current sequences (Figure 5A and Supplemental file 5).

**Figure 5.**
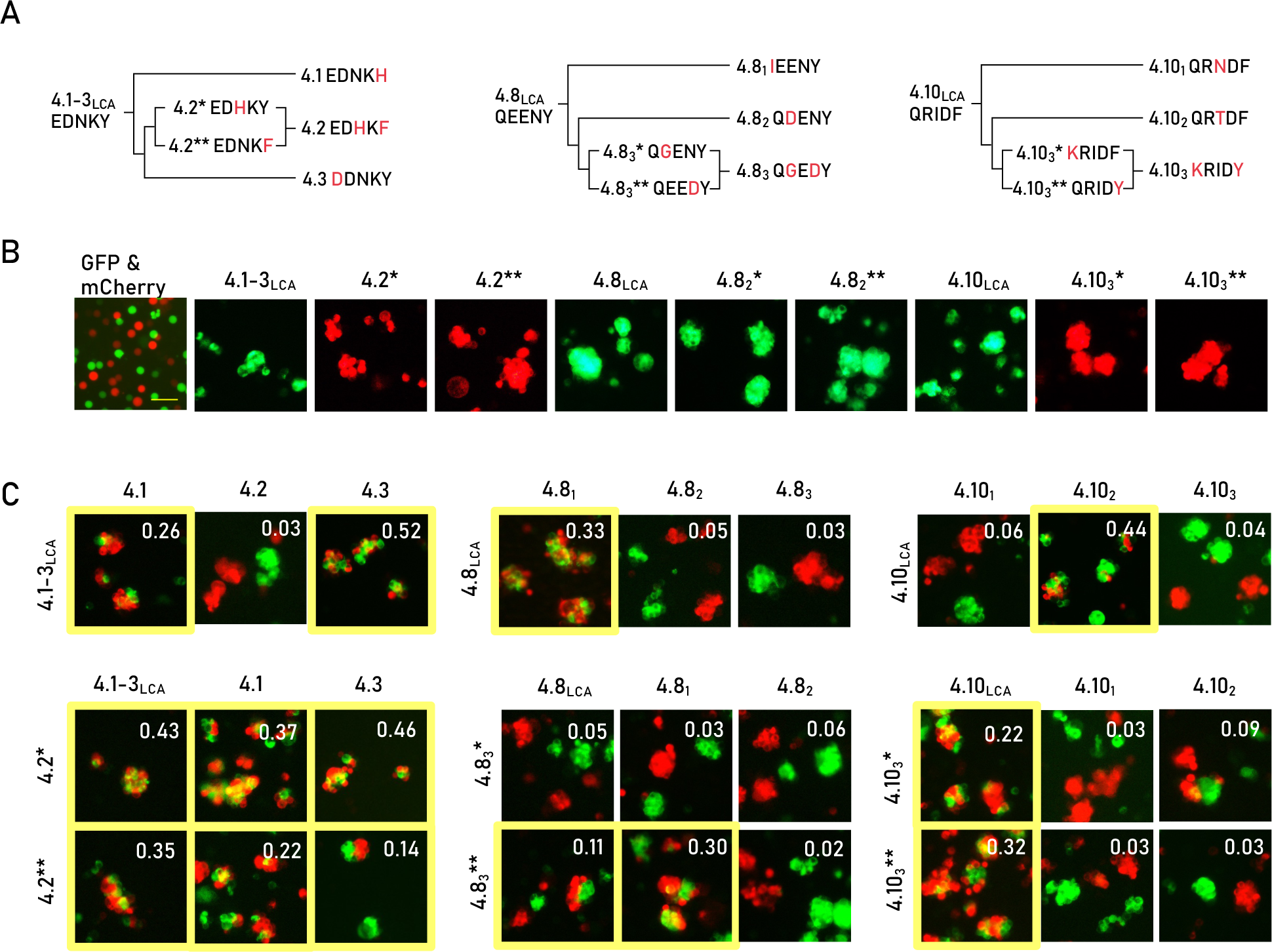
Tracing Evolutionary Paths of Binary Specificities of Dscam1. A) Predicted mutational pathways from ancestor to contemporary isoforms. The Ig2:Ig2 interface residues corresponding to sequence positions 107,109,111,112, and 114 are shown for each isoform. Exon duplications are indicated with diverging lines. Mutations are highlighted in red. B) Resurrected ancestral and intermediate proteins mediate cell aggregation, while the control cells expressing GFP and mCherry do not mediate aggregation. C) Binding preferences of resurrected and contemporary isoforms demonstrate heterophilic interactions in many cases (highlighted by the yellow boundary). The aggregate mixing score is presented in the right corner of each image. Scale 100 µm.

#### Resurrected ancestral proteins bind homophilically

We resurrected these ancestral interfaces by mutagenesis of extant interface residues and examined their ability to self-bind using the cell aggregation assay. We found that all three ancestors effectively mediated cell aggregation, demonstrating their ability to function as homophilic adhesion receptors (Figure 5B). Next, we compared the binding preferences of these ancestor sequences with their extant descendants. We observed that cells expressing the ancestral 4.8_LCA_ formed mixed aggregates exclusively with cells expressing the extant 4.8_1_ isoform, but not with 4.8_2_ and 4.8_3_. Similarly, 4.10_LCA_ was observed to interact solely with one of its current descendants, 4.10_1_, and not with the remaining two isoforms (Figure 5C). These results implicate that post-duplication, the ancestral 4.8 and 4.10 exons evolved into their respective current isoforms (4.8_1_ and 4.10_1_, respectively), while the other duplicates diverged and developed new binding specificities. Interestingly, cells expressing the 4.1-3_LCA_ ancestor recognized cells expressing either 4.3 or 4.1 extant isoforms, but not 4.2 expressing cells (Figure 5C, left). These results indicate that these extant proteins diverged via a subfunctionalization mechanism (McClune and Laub 2020), by which the ancestor protein binds to a wider range of partners (in this case, two distinct isoforms) compared to its descendants (which here bind strictly homophilically).

Next, we examined whether divergence of the Ig2 domain interface occurred through intermediate proteins that maintain homophilic cell adhesion or whether mutations in intermediate proteins disrupt self-binding. We observed that in all three studied examples, the ancestor sequence differs from two extant exons by a single residue and by two residues from the third extant exon. For example, the 4.1-3_LCA_ interface is composed of the following five residues: “EDNKY”. A single substitution from Tyrosine at position 114 to Histidine is sufficient to reach the interface of exon 4.1 (EDNKH). Similarly, a single substitution of the 4.1-3_LCA_ interface at position 107 from Glutamate to Aspartate would generate an exon 4.3 interface (DDNKY) (Figure 5A). Editing the 4.1-3_LCA_ interface to the interface of exon 4.2 (EDHKF) requires at least two mutations, N111H and Y114F, with two possible intermediate interfaces, EDHKY and EDNKF, depending on the mutation order. Using similar logic, we identified interface intermediates from the 4.10_LCA_ and 4.8_LCA_ to 4.10_3_ and 4.8_3_ respectively (Figure 5A).

We then examined self-aggregation abilities of cells expressing each intermediate isoform, with the aim of testing the self-binding capabilities of intermediate states. Surprisingly, we observed that all intermediate proteins engaged in homophilic interactions, despite having interfaces that appeared incompatible for such interactions (Figure 5B). For example, within the homodimer interface of one 4.8 intermediate isoform, two negatively charged Aspartate residues are positioned in proximity upon dimerization, where they could potentially lead to electrostatic repulsion (Figure 6). These types of incompatibilities are thought to be central in preventing unwanted cross-talk between different Dscam1 isoforms (Figure 4) (Sawaya, et al. 2008). To assess changes in the binding affinity (ΔΔG_bind_) of this intermediate, we used two computational methods, FoldX and SSIPe (Schymkowitz, et al. 2005; Huang, et al. 2020). Both methods predicted that the N112D mutation would significantly destabilize homophilic interactions, with ΔΔG_bind_ exceeding 2 kcal/mol. These predictions indicate that negative constraints weaken self-interaction of the intermediate isoforms. Yet, our cell aggregation results show these constraints do not entirely disrupt adhesive functionality.

**Figure 6.**
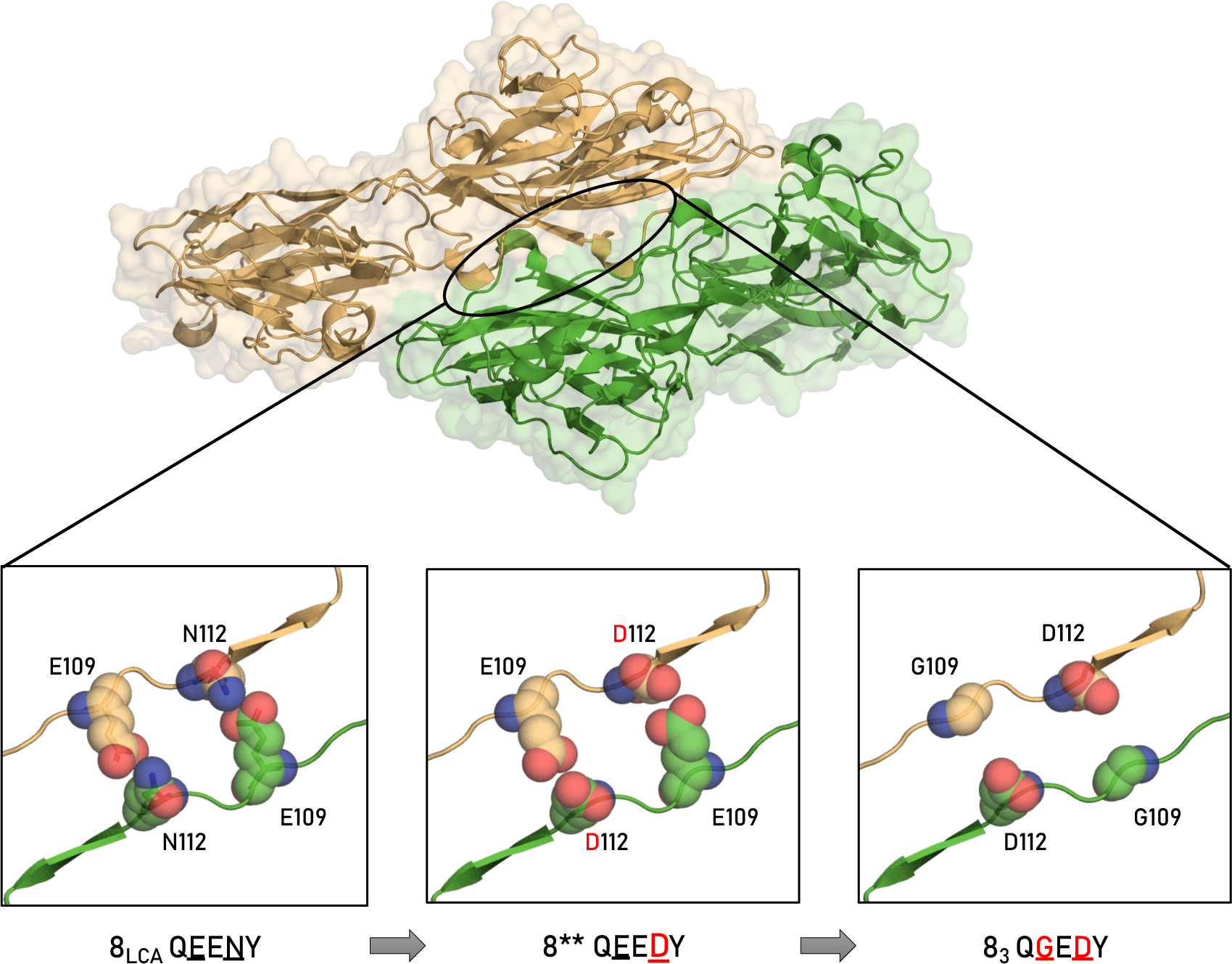
Dscam1 exon 8_3_ mutational path. The top image presents the Alphafold structural model of the Ig1-Ig4 dimer structure with Ig2 encoded by exon 4.8 ancestor and highlighted by a black oval. At the bottom, there are three close-up views of the structural model of the homophilic Ig2:Ig2 interface for 4.8 last common ancestor (left), 4.8 intermediate isoform (middle), and 4.8_3_ current isoform (right). Mutated residues are shown with Van der Waals spheres and colored based on chain origin and atom type (nitrogen in blue and oxygen in red). Ig2:Ig2 interface residues are noted at the bottom.

#### Resurrected ancestral proteins bind heterophilically with contemporary isoforms

We then tested whether non-specific binding could occur between ancestral, intermediate, and extant isoforms. We found that both alternative intermediate proteins that may lead to exon 4.2 formed mixed aggregates with extant isoforms 4.1, 4.3 and their ancestor, demonstrating heterophilic binding specificities (Figure 5D left). Thus, both mutations leading to exon 4.2 are necessary to prevent cross-interactions with closely related paralogs, generating strict homophilic binding. One of the intermediate proteins leading to 4.8_3_ recognized both the ancestor and one paralog extant sequence (4.8_1_). Finally, both intermediate proteins lead to 4.10_3_ promiscuously bound to the ancestor, with one intermediate binding weakly to one of the extant paralogs (4.10_2_) (Figure 5D). Overall, of the six intermediate proteins we tested, five exhibit promiscuous cross-interactions, demonstrating a gradual transition in specificity. Our experimental observations also explain the continued evolution of intermediate Dscam1 isoforms reconstructed here, as they generally exhibit non-specific cross-interactions with other isoforms.

## Discussion

This study examined the evolution of Dscam1 isoforms binding specificities, with a focus on the exon 4 cluster. This exon encodes the second immunoglobulin domain (Ig2), one of three domains involved in Dscam1 homophilic binding. We utilized a phylogenetic analysis, ancestral sequence reconstruction, and cell aggregation experiments and revealed several key insights: We identified relatively recent Dscam1 duplications that evolve strict homophilic binding in mosquitos. We also observed that Dscam1 proteins have maintained their fundamental functionality throughout their evolutionary trajectory, as both ancestral and evolutionary intermediate proteins mediate homophilic cell recognition. Finally, we discovered that in contrast to extant Dscam1 proteins that interact only homophilically, ancestral and intermediate proteins exhibit promiscuous interactions and are able to engage in both homophilic and heterophilic binding.

### Conservation of Dscam1 exon 4 cluster

With the goal of tracking evolutionary trajectories of the Ig2 dimer interface, our initial step involved a comprehensive search for current exon 4 sequences. Similar to past findings, we observed that exon 4 duplications are relatively conserved (Graveley, et al. 2004; Lee, et al. 2010; Armitage, et al. 2012), comprising nine invariant exons (Lee, et al. 2010). In addition to the nine conserved exons, we identified other exon 4 duplications across various insect lineages (Figure 2). These duplications expand the Dscam1 isoform repertoire by diversifying the Ig2 dimer interface sequences, albeit to a lesser extent than the significant expansions observed in exons 6 and 9. In all these instances, a single ortholog can consistently be identified through the conserved Ig2 dimer interface, while the other duplicated exons undergo non-synonymous substitutions, consistent with Ohno’s evolutionary model (Ohno 1970).

The conservation of most exon 4 variants could possibly be attributed to an unknown functional role these exon variants might have. Alternatively, it is possible that this conservation could be attributed to inherent constraints imposed by the small Ig2 dimer interface, comprising of only five residues.

### Strict homophilic binding in Dscam1 isoforms

To our knowledge, studies into the binding preferences of insect Dscam1 had previously been confined to *Drosophila melanogaster* isoforms. However, both the current and previous studies identified duplication events outside of the Drosophila lineage for which there yet to have been experimental investigations of their binding preferences (Graveley, et al. 2004; Lee, et al. 2010; Armitage, et al. 2012). Our findings reveal that even relatively recent duplications, occurring subsequent to the divergence of mosquitoes from flies, have evolved into isoforms exhibiting strict homophilic binding preferences (Figure 4ii & iii). Overall, these findings underscore the remarkable ability of Dscam1 proteins to generate self-binding domains via exon duplication and sequence divergence. These results also highlight the evolutionary pressure to generate isoforms that can accurately differentiate self from non-self interactions, which is central in neuronal patterning.

A previous study using ELISA showed that a small subset of exons, including exons 4.1 and 4.3, engage in both homophilic and significantly lower-affinity promiscuous heterophilic interactions (Wojtowicz, et al. 2007), in contrast to our findings. Interestingly, the authors themselves have observed such discrepancies in results between ELISA and cell aggregation assays. This difference in findings likely stems from the fundamental methodological differences between the assays. ELISA assays are more quantitative and sensitive to slight variations in protein binding affinities. On the other hand, cell aggregation assays offer an advantage by facilitating binding interactions within the context of native cellular membranes, potentially providing a more physiologically relevant measure of adhesion specificity.

### Ancient DSCAM1 proteins bind promiscuously

Understanding the emergence of new binding specificities is a central question in biochemistry and evolution. Our study investigates the evolutionary trajectories of insect Dscam1 exon 4 by focusing on recent duplications in flies and mosquitoes to reconstruct ancestral isoforms. When reconstructing the evolutionary paths from ancestral proteins to current isoforms, we found that as little as one or two residue changes were sufficient to alter the ancestral binding specificity. We reconstructed intermediate isoforms by examining the shortest mutational paths and specifically testing the impact of the two mutations that altered binding specificity. While the reconstruction of intermediate isoforms did not rely on statistical data, our conclusion that these intermediate isoforms retain self-binding capability is likely to be robust to this uncertainty. This is because both identified mutations are likely to be responsible for insolating the isoform from cross heterophilic interactions with its closely related paralogs. By introducing one mutation at a time, we test the maximal impact of these alterations on interface stability without allowing for additional bridging mutations. Therefore, although the actual evolutionary trajectory may have been more complex, our analysis of the most extreme mutations that could impact binding specificity on this path strongly supports the validity of our conclusion.

We discovered that across various evolutionary trajectories, Dscam1 isoforms consistently maintained their self-binding capabilities. In addition to self-binding, several evolutionary intermediate isoforms demonstrated promiscuous heterophilic interactions. These results were particularly surprising considering the necessity for Dscam1 paralogs to achieve highly-precise, strict homophilic interaction specificities for their role in neuronal self-avoidance. Previous studies have shown that Dscam1 isoforms evolve interfaces compatible with only self-interactions, while interfaces generated by heterophilic interactions contain non-complementary electrostatic charges and shapes, thereby preventing heterophilic interactions (Sawaya, et al. 2008).

Given these insights, one might expect intermediate isoforms to possess a non-complementary interface that would hinder homophilic binding rather than the interface that allows for cross-interactions (Figure 6, middle). While an isoform with abolished homophilic binding would not mediate neuronal self-recognition and patterning, the potential fitness penalty for a dysfunctional isoform may be mitigated by the presence of up to 50 randomly expressed isoforms in each neuron (Neves, et al. 2004; Zhan, et al. 2004). Our study challenges these expectations by demonstrating that, in fact, intermediate Dscam1 isoforms retain self-binding through their evolutionary development.

A possible explanation for this unexpected observation lies in the high degree of conservation of the exon 4 cluster, suggesting a specialized functional role, potentially imposing an additional evolutionary constraint. It would therefore be interesting to study the evolution of exon 6 or exon 9 clusters, which have been shown to be non-conserved and therefore could potentially exhibit functional disruptive mutations.

In summary, our study into the evolutionary history of Dscam1 exon 4 across various insect lineages provides insights into the mechanisms by which cell adhesion proteins evolve distinct binding specificities crucial for the regulation of complex cellular processes. Mutations leading to promiscuous binding have previously been documented in other protein systems evolved under selection against cross-talk, including the toxin-antitoxin systems, hormone receptors and enzymes (Voordeckers, et al. 2012; Aakre, et al. 2015; Devamani, et al. 2016; Siddiq, et al. 2017; Lite, et al. 2020). Our findings suggest parallel phenomenon in adhesion proteins, where promiscuous intermediates constitute a likely evolutionary step toward achieving highly precise binding specificity.

## Materials and Methods

### Dscam1 exon 4 tree construction

The translated sequence coded by exon 4 of Drosophila melanogaster Dscam1 transcript variant BE (isoform 7.9.30, NCBI annotation NM001043041) was used as the query for identifying exon 4 duplications in other insect genomes. The tblastN algorithm was used with the default parameters against the RefSeq Genome Database in the NCBI portal (Johnson, et al. 2008; Coordinators 2016; O’Leary, et al. 2016). The search set was limited to Coleoptera (taxid:7041), Diptera (taxid:7147), Hymenoptera (taxid:7399) and Lepidoptera (taxid:7088). Each order was searched separately due to the dataset favorable bias towards Drosophila species sequences, which led to other species sequences being omitted from the search results. The search resulted in 962 sequences from 83 species (see supplemental file 1). From the 83 species, 37 are Drosophilidae species with highly similar (>95% identity) exon 4 duplications to *Drosophila melanogaster*. To avoid over-representation biases, we included only the 12 *Drosophila melanogaster* exon 4 duplications in further analyses (see Supplemental file 1 for sequence list).

518 high-confidence exon 4 sequences retrieved in this search were clustered by cd-hit-v4.8.1-2019-0228 (Li and Godzik 2006; Fu, et al. 2012) using 95% threshold, to remove identical sequences. The 266 representative sequences were aligned with an outgroup sequence, a translated exon 4 of *Ixodes scapularis* (deer tick) Dscam2 (NCBI accession XP_042144217). *I. scapularis* was previously used as an outgroup organism for Dscam1 insect alignments (Armitage, et al. 2012). Alignment was performed by MAFFT v7.490 plugin in Geneious Prime® 2020.2.4 (Katoh, et al. 2002; Katoh and Standley 2013; (https://www.geneious.com). 2020), using the progressive method FFT-NS-2 algorithm, legacy gap penalty and default settings (see Supplemental file 2). Phylogenetic tree was constructed using FastTree 2.1.11 plugin in Geneious Prime® 2020.2.4 (Price, et al. 2009; (https://www.geneious.com). 2020) with WAG 2001 amino acid substitution model. The FastTree tree was rooted using Geneious Prime® 2020.2.4 branch rooting feature (see Supplemental file 3). Sequence similarity of the five Ig2 dimer interface positions (107, 109, 111, 112 and 114) was calculated using the BLOSUM62 matrix.

### Ancestry sequence reconstruction

Based on the rooted FastTree, ancestor sequences were predicted using two independent ancestral sequence reconstruction programs-PaMLX 1.3.1 CodeML (Yang 2007; Xu and Yang 2013) and GRASP (Ross, et al. 2022). For detailed accuracy estimation see supplementary file 2.

### Cloning

Dscam1 coding sequence was reconstructed by cloning isoform 7.27.25 ECD (Ig1-9 and the first FNIII domain) and the remaining FNIII domains, transmembrane and cytoplasmic domains from clone RE54695 (barcode 9407 from the Drosophila Genomics Resource Center, NIH Grant 2P40OD010949). Both sequences were cloned using Gibson assembly (NEBuilder® HiFi DNA Assembly Cloning Kit E5520S) into a modified lenti virus (pLV) transfer plasmid.

Site-directed mutagenesis was performed on the sequence encoding to the Ig2 dimer interface in pCMVi-Dscam1_7.27.25-AP using QuikChange® Lightning Site-Directed Mutagenesis Kit 210518. Mutagenesis primers were designed with the QuikChange® primer design program. Positions 1-3011 of the mutant construct was amplified and cloned into plv-Dscam1_7.27.25(Δ1-3011)-GFP and plv-Dscam1_7.27.25 (Δ1-3011)-mCherry amplicons using Gibson assembly.

### Cell aggregation assay

FreeStyle™ 293-F cells (Thermo Fisher R79007) were separately transfected using PEI MAX® - linear polyethylenimine hydrochloride (MW 40,000, 49553-93-7) as follows: 1 million cells per ml were grown in FreeStyle™ 293 Expression Medium at 6 wells non-treated plates (SPL #32006), 2 ml per well at 37°C, 8% CO_2_, 135 rpm. 1.25 µg plasmid DNA was mixed in Opti-MEM I reduced serum media (Thermo Fisher 11058021) to a final volume of 62.5 µl. 3.12 µl PEI was added with Opti-MEM I media to a final volume of 62.5 µl. Both mixtures were incubated at room temperature for 15 minutes. The PEI mix was then added to the DNA mix, immediately vortexed and incubated at room temperature for an additional 15 minutes, then added to the cells. 3 hours later 500 µl FreeStyle™ 293 Expression Medium was added to the cells. 1 ml of each reaction was mixed with 1 ml of the complementary reaction for testing binding preferences of desired isoforms. Three biological replications for each isoform combination were performed, with eight images acquired per replication using Eclipse Ts2 inverted microscope 48 hours post transfection.

### Aggregate mixing quantification

Figures 4 and 5 illustrate the aggregation or separation of cells, which was quantified using a custom Python script. This script determines the ratio of red and green cells in close proximity to each other. To achieve this, the script divides the images into a grid and examines each grid cell to identify both red and green HEK293F cells. Each grid cell covers an area of 24.32 µm², approximately equivalent to 1.5-2 cells in size. Hence, if both red and green HEK293F cells are found within the same grid cell, it indicates potential contact between them. The images shown in Figures 4 and 5 are cropped to depict representative aggregates. However, the quantification of mixing was carried out across the entire image area for all acquired images. The proportion displayed represents the average score obtained from three technical replications, with eight acquired images for each replication of each isoform combination. For further statistical details, refer to supplemental figure 1.

### Computational evaluation of binding affinities

Structural model of DSCAM1 8_LCA_ was prepared using alphafold2 (Jumper, et al. 2021; Mirdita, et al. 2022). The structure was refined using at least five rounds of the “RepairPDB” utility in foldX (Schymkowitz, et al. 2005). Mutations were generated using the “BuildModel” utility and analyzed using “AnalyseComplex” utility. Additionally we used the SSIPe server with default parameters for calculating the difference in binding energy upon mutation (Huang, et al. 2020).

## Acknowledgments

We thank Prof. Dinorah Morvinski from the department of biochemistry and molecular biology at Tel Aviv university for providing the lenti virus (pLV) transfer plasmid. We thank Dr. Karin Smorodinsky-Atias for her assistance during the experimental stages of this work. This work was supported by the Israel Science Foundation (1463/19 to R.R).

