## Supplementary material for "Following The Evolutionary Paths Of Highly Specific Homophilic Adhesion Proteins": Supplementry Figure 1.pdf

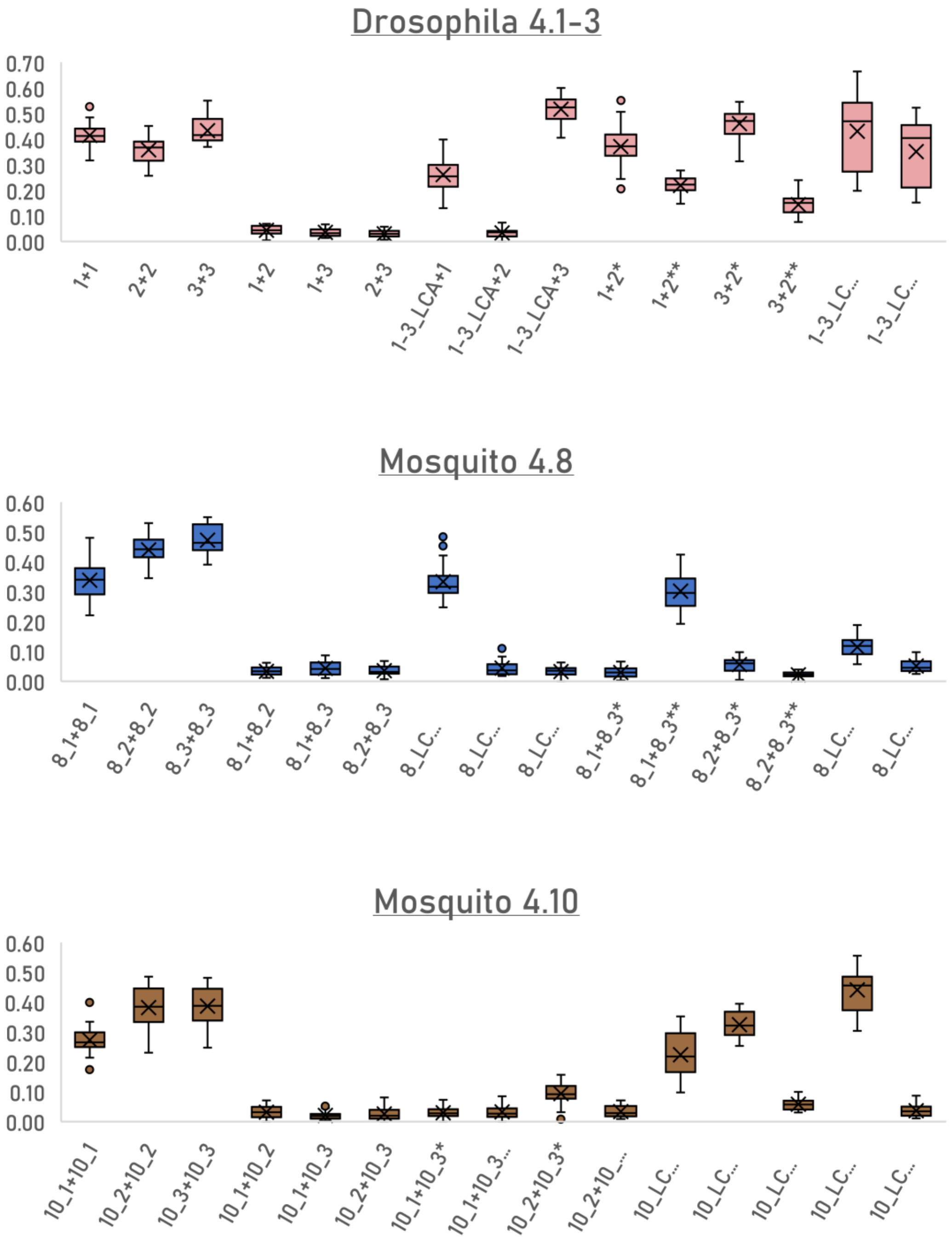


**Supplementry Figure 1.** (related to main Figures 4 and 5). Statistical analyses of the mixing scores calculated for co-aggregation assays. Mixing scores are assigned in automated image analyses (see Methods).
